# A theoretical framework for how ecological interactions between microbes affect mutant fitness

**DOI:** 10.64898/2026.05.22.727324

**Authors:** Justus Wilhelm Fink, Duhita G. Sant, Michael Manhart

**Author notes:** These authors contributed equally (Justus Wilhelm Fink, Duhita G. Sant).

## Abstract

The distribution of fitness effects (DFE) for spontaneous mutations characterizes both an organism’s evolutionary potential as well as its genomic functions. The DFE of a genome depends on the specific environment in which it is measured, and for microbes a major feature of their environment is the presence of interactions with other species, such as competing for or cross-feeding nutrients. Several recent studies have empirically measured how the DFE of one microbial species changes in the presence of interactions with other species. However, the underlying mechanisms by which this happens, and the statistical patterns they are expected to produce, are unknown. Here we classify two types of statistical changes in the DFE: global changes to the DFE, such as to its mean or variance, and idiosyncratic changes in the fitness of individual mutants, summarized by the correlation of mutant fitness between environments. We first show that both types of effects occur in empirically measured DFEs across a wide range of species and interactions; idiosyncratic effects appear to have a maximum limit and constrain the size of global effects. We then show that a minimal model of an ecological interaction (competition for a single resource) is sufficient to generate both types of effects. Finally, we extend this model to arbitrary quantitative traits to reveal two general mechanisms of how interactions alter the DFE: 1) interactions can globally change fitness by altering the community growth rate, and 2) interactions can idiosyncratically change fitness of individual mutants by altering relative selection on different traits affected by those mutations.

## INTRODUCTION

The distribution of fitness effects (DFE) — the set of fitness values for all spontaneous mutations available to a genome [1] — is a key input to predicting how a population will evolve [2–4] as well as an indicator of genomic functions [5]. While previous studies have attempted to measure or infer the DFE in many systems, especially microbes, a major limitation to using the DFE for evolutionary predictions is that it depends significantly on the environment [5–12]. For microbes, a major feature of their environments is the presence of other microbes that a focal species interacts with, such as by competing for [13–15] or cross-feeding nutrients [16–18]. These interactions can change over time, for example in the gut microbiome of people with certain diseases [19–22]. Several recent studies have found that ecological interactions between microbes can significantly change the DFE [23– 31] and consequent adaptation [32–37]. Since we cannot empirically measure a microbe’s DFE under all possible interactions, we need general principles for the DFE’s environmental dependence if we hope to make predictions in natural systems where environments vary over time and space.

Despite the clear evidence that interactions alter the DFE, exactly how and why they do so is unclear. In some species, mutualistic cross-feeding interactions tend to reduce the magnitude of mutant fitness [27– However, in other species, interactions have apparently random effects on mutant fitness [24, 25, 27, 31], and some interactions do not significantly change mutant fitness at all [30, 38]. A particular study by Martinson et al. [30] directly demonstrated this complexity for a single species: they found that while cross-feeding tended to change both the DFE mean and the fitness of many individual mutants (mainly rescuing deleterious ones) in *Salmonella enterica*, competition left the DFE of the same species largely unaffected. The abundance of recent empirical observations has created a need for theory and models to understand which results are surprising and which can be explained by simple principles. For example, should we expect competitive and cross-feeding interactions to change the DFE in different ways? How should those DFE changes depend on traits of the other species? These questions are important to understand how different types of ecological environments dictate evolutionary outcomes: an environment dominated by cross-feeding interactions (e.g., in the dysbiotic gut microbiome [22]) might induce DFEs with different evolutionary outcomes compared to an environment dominated by competition (e.g., in the healthy gut).

This article aims to build the foundation for a theory to interpret these experiments. We first introduce a statistical scheme for classifying as global or idiosyncratic the effects an interaction (or any environmental factor) can have on the DFE. We demonstrate the empirical prevalence of both effects across a wide range of species and interactions. We then test for these effects in a minimal model of resource competition, as arguably the simplest ecological interaction and one believed to dominate ecological dynamics in many natural microbial communities [13–15]. By simulating mutations as random perturbations in traits, we show how competitive interactions can alter the DFE both globally and idiosyncratically when the competitor species grows faster than the focal species, by increasing the community growth rate and changing relative selection on traits. Finally, we show how these results generalize to arbitrary ecological interactions and population dynamics using a quantitative traits model.

## RESULTS

### Ecological interactions can change mutant fitness globally or idiosyncratically

Inspired by previous work on epistasis [39] (gene-by-gene interactions), we introduce a statistical scheme for classifying how an ecological interaction changes the fitness of a mutant (gene-by-environment interactions). The effect of the ecological interaction is defined by how it changes the monoculture fitness 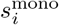 to become the coculture fitness 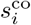 of a mutant *i*. The interaction can have a *global* effect that transforms the fitness of all mutants the same way, i.e., there is a single function *f* (*s*^mono^) = *s*^co^ that relates monoculture to coculture fitness (Supplementary Sec. S1). This means the interaction changes the DFE’s overall statistics, such as its mean, variance, or other moments [7]. For example, the interaction might shift the fitness of all mutants by a constant, thereby changing the mean of the DFE (Fig. 1A, top panel), or rescale each mutant’s fitness, thereby changing the variance (Fig. 1A, bottom panel). In contrast, the interaction can also have *idiosyncratic* effects that are unique to individual mutants, so that there is no single transformation *f* from monoculture to coculture fitness, but instead a distinct function 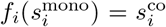 for each mutant *i* (Supplementary Sec. S1). Over the whole DFE, this is summarized by the correlation coefficient of fitness for individual mutants between the two environments: idiosyncratic effects make the correlation coefficient less than 1 (Fig. 1B).

**FIG. 1.**
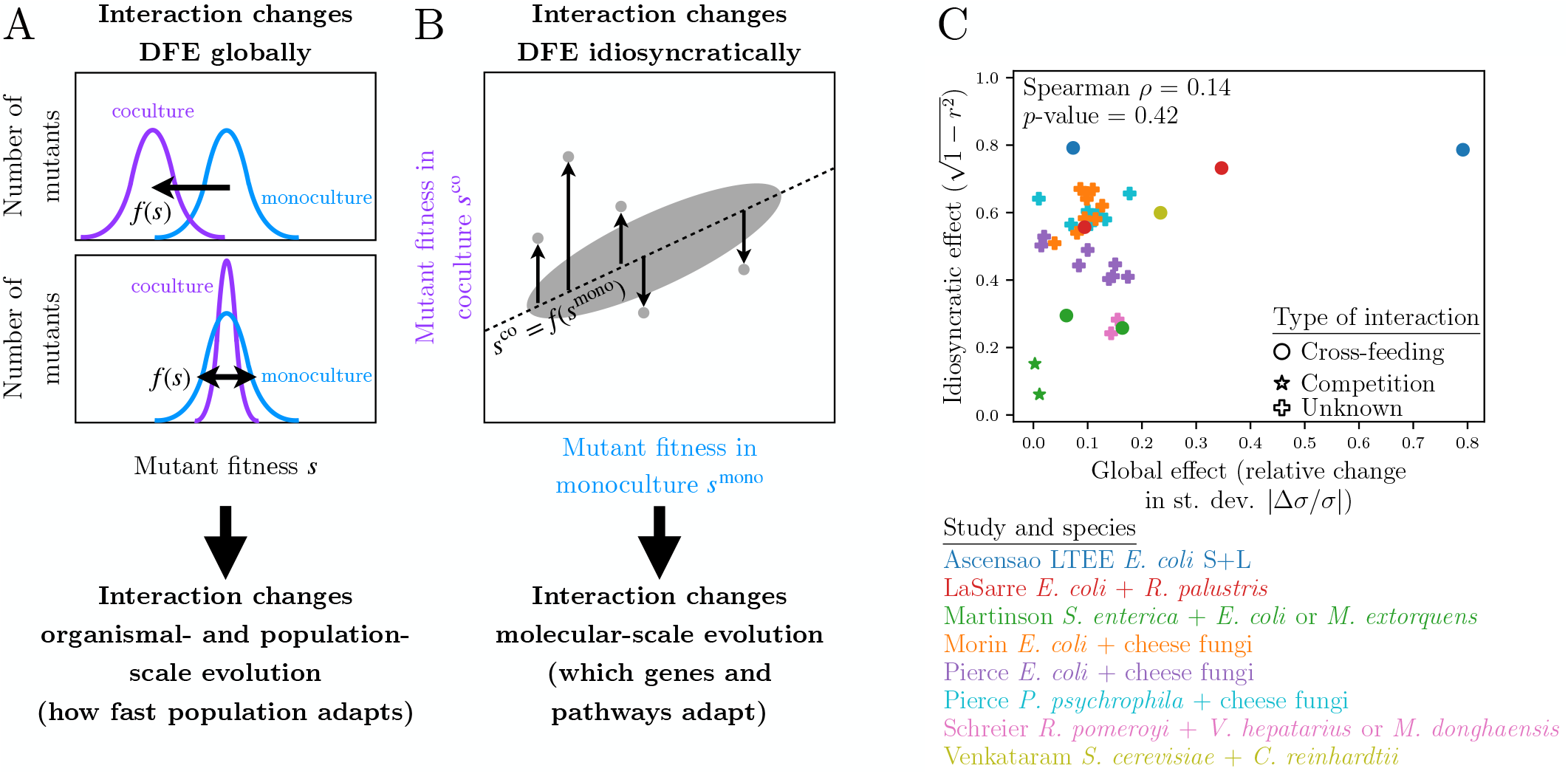
Possible statistical effects of an ecological interaction on the DFE. (A) Schematic DFEs with global effects of an interaction (*s*^co^ = *f* (*s*^mono^)), changing the mean (top panel) or variance (bottom panel) and altering consequent evolution at the organismal and population scales. (B) Schematic scatter plot of fitness in monoculture vs. coculture with idiosyncratic effects of an interaction. The dashed diagonal line represents the global effect, with vertical arrows representing idiosyncratic deviations from it (Supplementary Sec. S1) that alter consequent evolution at the molecular scale. (C) Comparison of global effects (measured as the relative change in the DFE standard deviation) and idiosyncratic effects (measured as 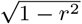, where *r* is the Pearson correlation coefficient) on the DFE from ecological interactions, across a range of published data sets (Methods, Supplementary Table S1).

Classifying the effect of an interaction on mutant fitness as global or idiosyncratic is useful for two reasons. First, these two types of effects are mathematically independent. Second, these two types of effects correspond to distinct biological consequences of the interaction on evolution (Fig. 1A,B). Global effects on mutant fitness mean the interaction alters evolution at the organismal and population scales [32, 33, 35]. For example, increasing the mean beneficial mutant fitness will increase the population’s adaptation rate [2]. Idiosyncratic effects, on the other hand, change evolution at the molecular scale: different genetic mutations may fix or be purged in the presence of the interaction compared to its absence [34, 36, 37].

To demonstrate the empirical existence of both effects, we collect 36 published data sets comparing an organism’s DFE in monoculture to its DFE in coculture with another strain or species (Methods, Supplementary Table S1, Supplementary Figs. S1 and S2). Figure 1C shows that these interactions generate both global and idiosyncratic effects, which span a wide range of magnitudes. Most interactions globally change the standard deviation by 10–20%, but a few interactions change it as little as 0.3% (e.g., *E. coli* or *M. extorquens* competing with *S. enterica* [30]), while others change it up to 80% (e.g., *E. coli* LTEE S strain cross-feeding with the L strain [29]). Idiosyncratic effects also span a wide range, from 6% (for *M. extorquens* competing with *S. enterica* [30]) to 80% (for either the LTEE L or S strains of *E. coli* LTEE cross-feeding [29]), with most being 40– 60%. Interestingly, three separate data points are clustered near ∼80% idiosyncratic effects (corresponding to a correlation coefficient of *r* = 0.6), which raises the question of whether there is a maximum limit of how much an interaction can change individual mutant fitness.

Global effects differ for some systems using different metrics (Supplementary Fig. S3); in particular, idiosyncratic effects correlate strongly with the global effect on the DFE mean (Supplementary Fig. S3A) but not with the global effect on the standard deviation (Fig. 1C). However, for all global effect metrics, they appear to be bound by the magnitude of idiosyncratic effects. That is, idiosyncratic effects set a maximum limit on the size of global effects, so that the interactions generating the smallest or largest global effects tend to correspond to the interactions generating the smallest or largest idiosyncratic effects. This also means that there are interactions with large idiosyncratic effects and small global effects, but not the other way around.

### A minimal model of mutant fitness under an ecological interaction

While empirical data demonstrates the existence of global and idiosyncratic effects of interactions on DFEs, it lacks a clear pattern in terms of the species or interaction, especially since so many interactions are of unknown type. We therefore need models to help understand the range of possibilities for how ecological interactions generate these effects. We start with a minimal model of an ecological interaction in microbes: competition for a single limiting resource (Fig. 2). While other interactions (cross-feeding [16–18], antagonism [40, 41], predation [42]) are also important in many microbial communities, we focus on resource competition as a test case since it is present in all ecosystems to some degree and is believed to dominate ecological dynamics in many natural microbial communities [13– As we will show, the model not only demonstrates that minimal assumptions about the interaction and population dynamics are necessary to generate both types of effects on mutant fitness, but it also reveals the general mechanisms by which any interaction (not just competition) alters mutant fitness.

**FIG. 2.**
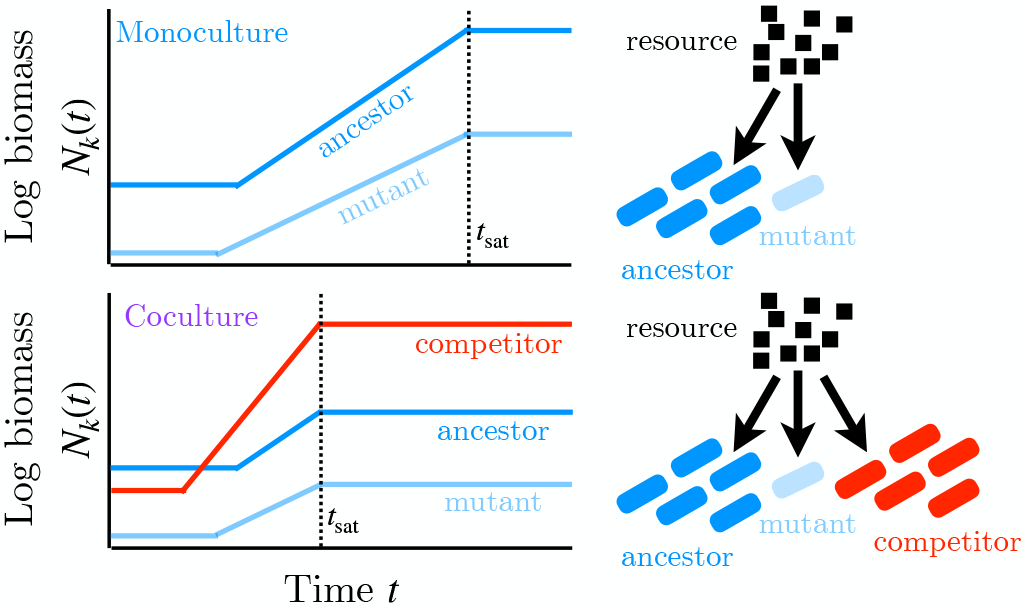
Minimal model of a competitive interaction between species. Biomass of genotypes grows in three phases (lag, exponential growth, stationary). Their growth saturates at time *t*_sat_ when the single limiting resource is exhausted (Methods).

Let the biomass concentration of each genotype *k* be *N*_*k*_(*t*), where *k* refers to one of three genotypes: an ancestral genotype (*k* = anc), a mutant of that ancestor (*k* = mut), or a competitor species (*k* = com). We assume the community grows in a batch culture, such that it receives an initial pulse of resources at *t* = 0 and grows until time *t* = *t*_sat_ when resources are depleted and biomass saturates (Fig. 2, Methods). We focus on batch cultures since laboratory measurements of microbial fitness almost always use batch cultures [24, 25, 27–31, 43], and they serve as a model for natural environments where biomass growth is fast compared to the frequency of resource pulses [44].

We use a minimal model of population dynamics, investigated in previous work [45, 46], that captures the basic sigmoidal shape of a microbial batch growth curve (Fig. 2, Methods): genotype *k*’s growth rate is zero until the lag time *λ*_*k*_, after which biomass starts growing exponentially at constant rate *g*_*k*_ and producing *Y*_*k*_ new units of biomass per unit resource (the yield) until the resource concentration reaches zero, at which time (*t* = *t*_sat_) growth immediately halts for all genotypes (Fig. 2). One could carry out a similar analysis as we do here with other common models of competition, such as the same consumer-resource model but with a Monod dependence for the growth rate [47, 48], or a Lotka-Volterra model [49]. However, the lag-growth model we use here is more mathematically tractable and easier to interpret in terms of population dynamics mechanisms than both of those other models are, and hence we believe it pro-vides a better demonstration of the general principles by which ecological interactions affect mutant fitness.

To calculate the DFE in this model, we simulate a large number of mutants as being random perturbations of their ancestor’s lag time and maximum growth rate (Methods). For each mutant we calculate its fitness relative to the ancestor as [43, 45, 46]

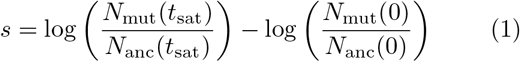

in a monoculture with its ancestor as well as in a coculture with the ancestor and a competitor species (Methods).

### Interactions globally rescale mutant fitness by changing community growth rate

To demonstrate the global effects of a competitive interaction on mutant fitness in our model, Fig. 3A shows three example DFEs: one in monoculture (no competitor), one in coculture with a slow competitor (*g*_com_ < *g*_anc_), and one in coculture with a fast competitor (*g*_com_ > *g*_anc_). Competition does not shift the DFE mean but does rescale the width of the DFE, with a faster competitor making the DFE much narrower (weaker selection), but a slower competitor making the DFE only slightly wider (stronger selection). To test this systematically, we scan traits of the competitor and measure the standard deviation of the DFE as its width. This shows competitors that are either faster (larger growth rate, Fig. 3B) or less efficient (lower yield, Fig. 3B) can significantly shrink the DFE width, whereas competitors with the opposite trait patterns increase the width only slightly. Competitor lag time has only modest global effects on the DFE (Supplementary Fig. S4A).

**FIG. 3.**
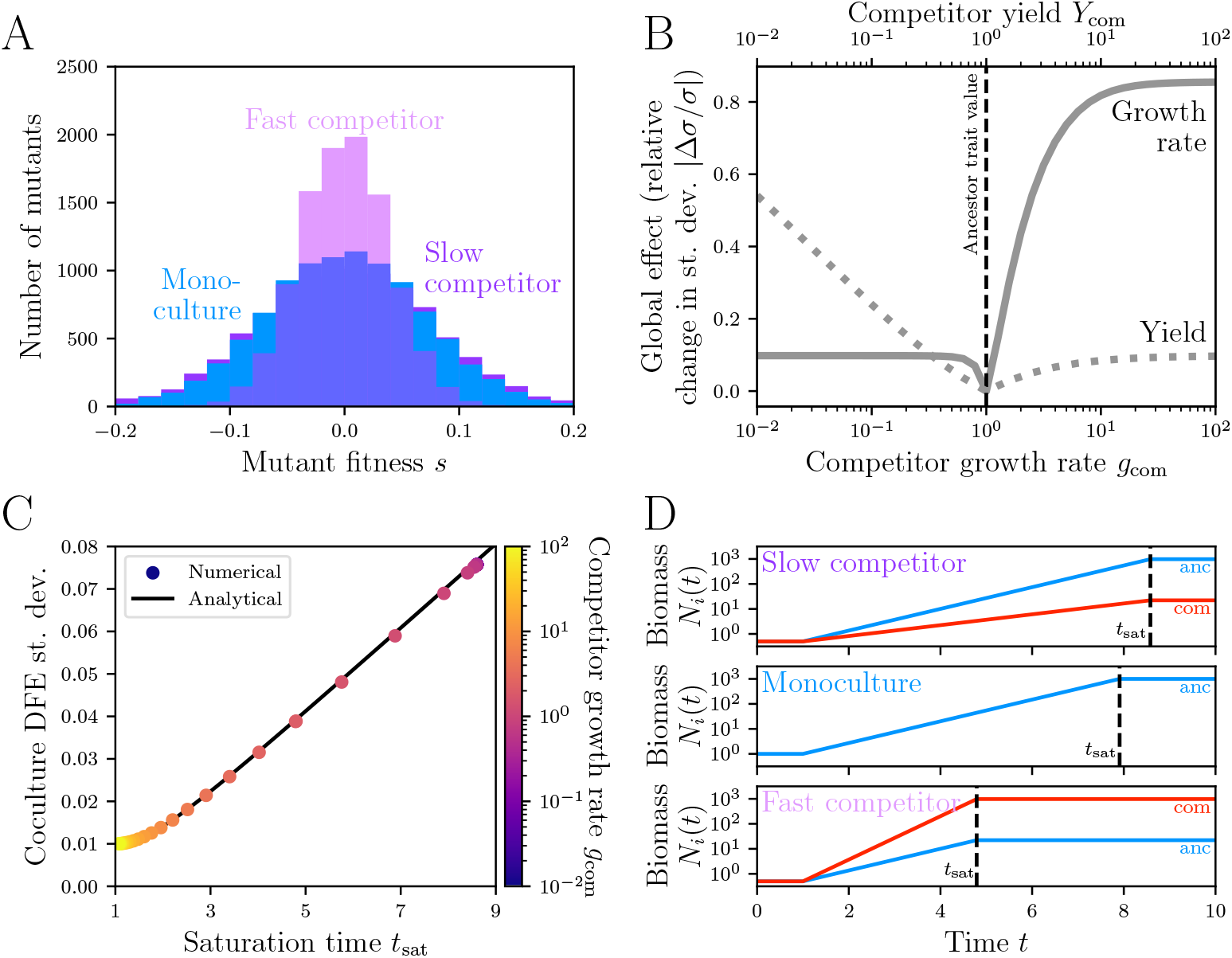
Cocultures with fast or inefficient competitors globally reduce DFE width. (A) Example DFEs from a monoculture without a competitor (blue), a coculture with a slow competitor (dark purple, *g*_com_ = 0.5), and a coculture with a fast competitor (light purple, *g*_com_ = 2). (B) Global effect of competition on the DFE (magnitude of relative change in standard deviation) as a function of competitor growth rate *g*_com_ (solid line, bottom axis) and competitor yield *Y*_com_ (dotted line, top axis). (C) Dependence of DFE standard deviation on saturation time *t*_sat_, across cocultures with varying competitor growth rates *g*_com_. Dots are numerical calculations (colored according to *g*_com_, same data as in panel B). The solid black line is the analytical calculation (Eq. 2). (D) Growth curves and saturation time *t*_sat_ of the ancestor (blue) and competitor (red) for the same three scenarios as in panel A.

In Supplementary Sec. S2 we analytically calculate the mean and variance of the DFE in the model. Under the assumptions of our mutation simulations (Methods), the DFE mean is always zero and the DFE variance is:

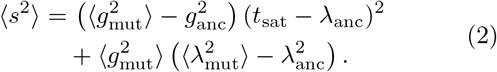

As this shows, the effect of the interaction on fitness is mediated solely by the saturation time *t*_sat_ when resources are depleted; this is a consequence of the fundamental assumption of consumer-resource models, that resource availability mediates all interactions between species (Methods). Thus the competitor globally changes mutant fitness in our model by changing *t*_sat_, which we can think of as the reciprocal of the community’s overall growth rate: a shorter *t*_sat_ means the community as a whole grows faster. Figure 3C shows how the DFE width depends on *t*_sat_ in numerical calculations of the model, which exactly matches the analytical expression in Eq. 2 (see also Supplementary Sec. S3 and Fig. S4B).

The centrality of the community growth rate also explains why slower and faster (or, less efficient and more efficient) competitors have asymmetric effects on the DFE width (Fig. 3A,B). The saturation time depends strongly on whichever genotype in the community is fastest or least efficient [45, 46, 50] (Supplementary Sec. S4). So adding a slower or more efficient competitor increases *t*_sat_ only a little (Fig. 3D, compare top and middle panels), while adding a faster or less efficient competitor decreases *t*_sat_ a lot (Fig. 3D, compare middle and bottom panels).

### Interactions change mutant fitness idiosyncratically when there is pleiotropy between traits

We now turn to addressing whether the competitive interaction of our model idiosyncratically changes the fitness of individual mutants (Fig. 1B). In Fig. 4A we show scatter plots of individual mutant fitness with and without an interaction. As with global effects on the DFE width, we see that decorrelating the DFE occurs with a fast competitor but not with a slow competitor (Fig. 4B). Unlike the global effects, though, the idiosyncratic effects depends crucially on the mutants being pleiotropic, i.e., affecting both growth rate and lag time simultaneously (Fig. 4B, Methods).

**FIG. 4.**
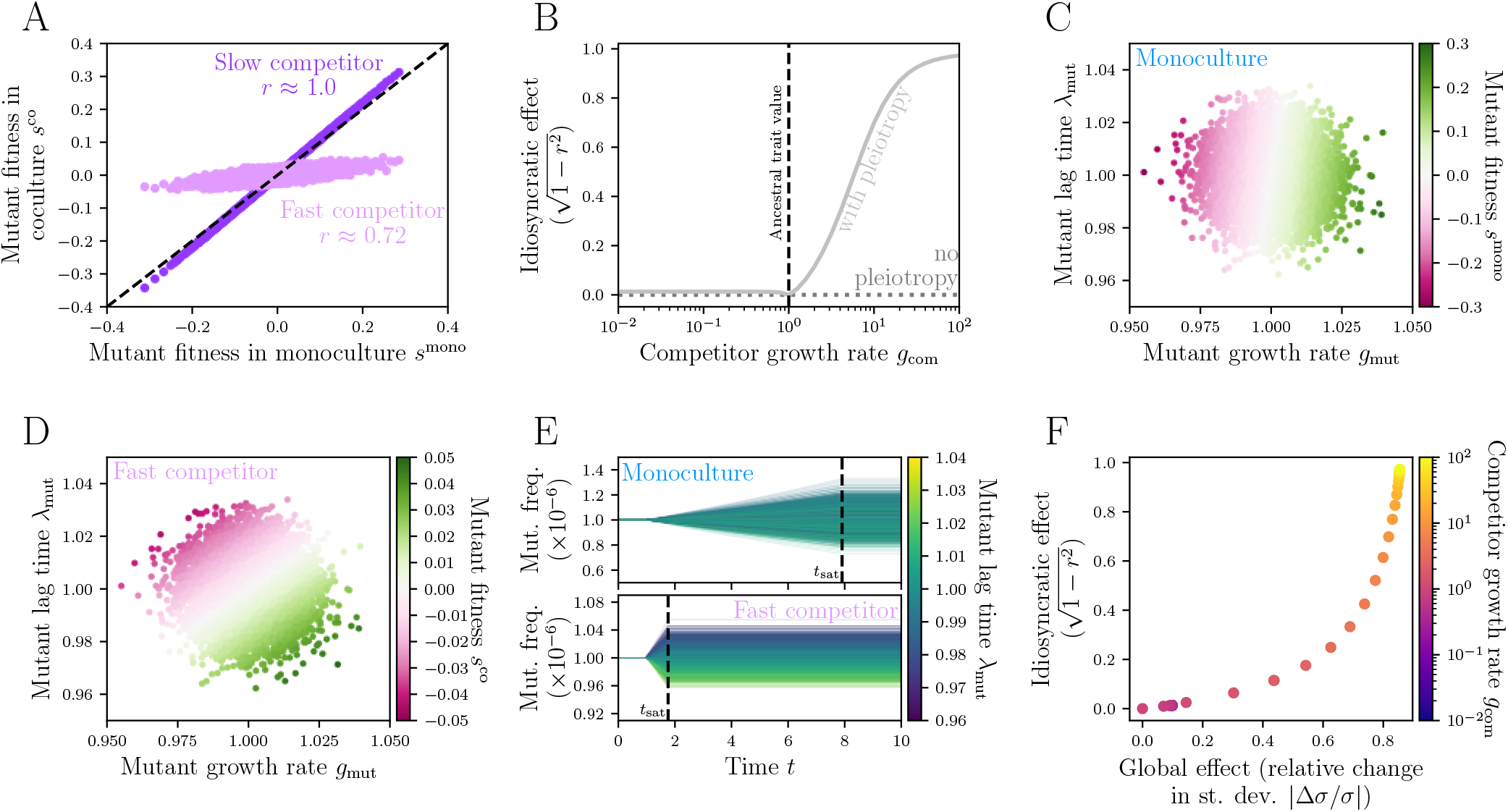
Cocultures with fast competitors idiosyncratically change mutant fitness when there is pleiotropy. (A) Scatter plot of fitness values for individual mutants in monoculture (*s*^mono^) and in coculture (*s*^co^), with either a slow (dark purple, *g*_com_ = 0.1) or a fast (light purple, *g*_com_ = 10) competitor. (B) Idiosyncratic effect (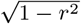, where *r* is the Pearson correlation coefficient) of the competitive interaction, as a function of competitor growth rate *g*_com_. Dotted dark gray line is for mutants without pleiotropy (mutations affect growth rate only, no variation in lag times), and the light gray line is for pleiotropic mutants (affecting both growth rate and lag time). (C) Scatter plot of mutant growth rates *g*_mut_ and lag times *λ*_mut_ colored according to their fitness in monoculture *s*^mono^. (D) Same as a panel C but colored according to fitness in coculture *s*^co^ with a fast competitor (*g*_com_ = 10). (E) Frequency of mutants (relative to ancestor biomass) over time *t* in monoculture (top panel) and in coculture with a fast competitor (bottom panel, *g*_com_ = 10). Colors of mutant trajectories in dicate their lag times *λ*_mut_. (F) Global effects (measured as relative change in standard deviation) vs. idiosyncratic effects (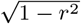, where *r* is the Pearson correlation coefficient) of competitors on the DFE. Dots are numerical calculations colored according to the competitor growth rate *g*_com_ (compare to empirical data in Fig. 1C).

We can see this mathematically by calculating the covariance between fitness *s*^mono^ in the monoculture (without competition) and fitness *s*^co^ in the coculture (with competition):

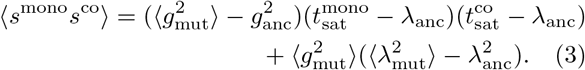

When there is no pleiotropy, e.g., mutants only affect growth rate and not lag time 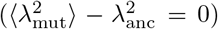, the term in Eq. 3 proportional to the lag time variance drops out, and the remaining term is simply the product of standard deviations for *s*^mono^ and *s*^co^ (using Eq. 2 with no lag time variance); hence there is a perfect correlation in the case without pleiotropy (Fig. 4B). (In the other pleiotropic case where mutations affect lag time but not growth rate, the interaction does not change fitness at all because fitness is independent of *t*_sat_.) Otherwise, the correlation coefficient is less than 1 in general and the interaction has idiosyncratic effects on individual mutants.

A conceptual way to think about idiosyncratic effects is that a fast competitor, by significantly reducing *t*_sat_, changes the relative selection on lag time vs. growth rate: with a slow competitor or no competitor (monoculture), a mutant’s fitness depends mostly on its growth rate (Fig. 4C), while with a fast competitor, mutant fitness depends more on its lag time (Fig. 4D; compare both cases in Fig. 4C,D with systematic scan in Supplementary Fig. S5). This is because *t*_sat_ is so short with a fast competitor that only the mutants with short lag times start growing and increase their frequencies (Fig. 4E). The asymmetric roles of faster and slower competitors on idiosyncratic effects (Fig. 4A,B) is caused by the same mechanism as for global effects (Fig. 3D, Supplementary Sec. S4), as both are mediated by *t*_sat_.

Altogether the model shows that faster or less efficient competitors can cause both global and idiosyncratic effects on the fitness of pleiotropic mutants. Analogous to our analysis of the empirical data sets (Fig. 1C and Supplementary Fig. S3), we can look at the variation of both types of effects across competitive interactions in the model (Fig. 4F and Supplementary Fig. S6). Both effects increase as competitors get faster, but we see a different sensitivity than in the empirical data. While in the model global effects are more sensitive to interactions than are idiosyncratic effects, the empirical data seems to show the opposite, that interactions more readily generate idiosyncratic rather than global effects.

### A quantitative traits model demonstrates how ecological interactions change mutant fitness in general

While the foregoing results demonstrate the potential for an ecological interaction to change mutant fitness in a simple model of resource competition, we can show how those results generalize to interactions and population dynamics of arbitrary complexity using a quantitative traits model. Let the growth of the ancestral species depend on a set of quantitative traits {*z*_𝓁_}; in the competition model, these traits were the lag time *λ* and growth rate *g*. Assuming the mutation only makes small relative perturbations to these traits, we can approximate its fitness as (using Eq. 1, Supplementary Sec. S5):

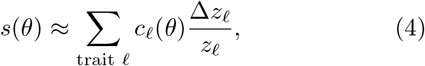

where the selection gradient *c*_𝓁_ (*θ*) on trait 𝓁 [51] is a function of a parameter *θ* that characterizes the ecological interaction (e.g., presence and traits of an interacting species). Any traits *z*_𝓁_whose effects accrue per generation [52] will have selection gradients proportional to the community’s batch growth time scale, which should be inversely proportional to the growth rate of the whole community (Supplementary Sec. S5):

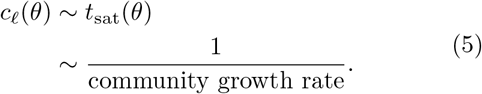

To generalize our previous results on how an interaction globally changes the DFE, we show how the DFE moments depend on the interaction parameter *θ*. The mean of the DFE is

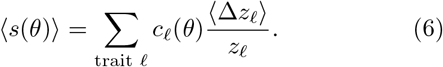

Assuming the interaction changes at least one of the selection gradients *c*_𝓁_ (*θ*) (e.g., by changing the community growth rate as in our competition model), then the interaction will also generally change the DFE mean if the average mutation trait effects ⟨Δ*z*_𝓁_⟩ are not zero for all traits 𝓁 (unlike the assumption in the competition model; Methods). We can similarly calculate the variance (compare to Eq. 2):

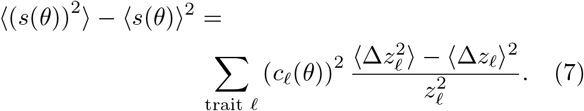

Under the assumption that selection gradients are inversely proportional to the community growth rate (Eq. 5), the mean and variance expressions (Eqs. 6 and 7) imply that a general global effect of an interaction is to rescale terms in all DFE moments by the community growth rate (Fig. 5A), just as we observed in our competition model (Fig. 3C).

**FIG. 5.**
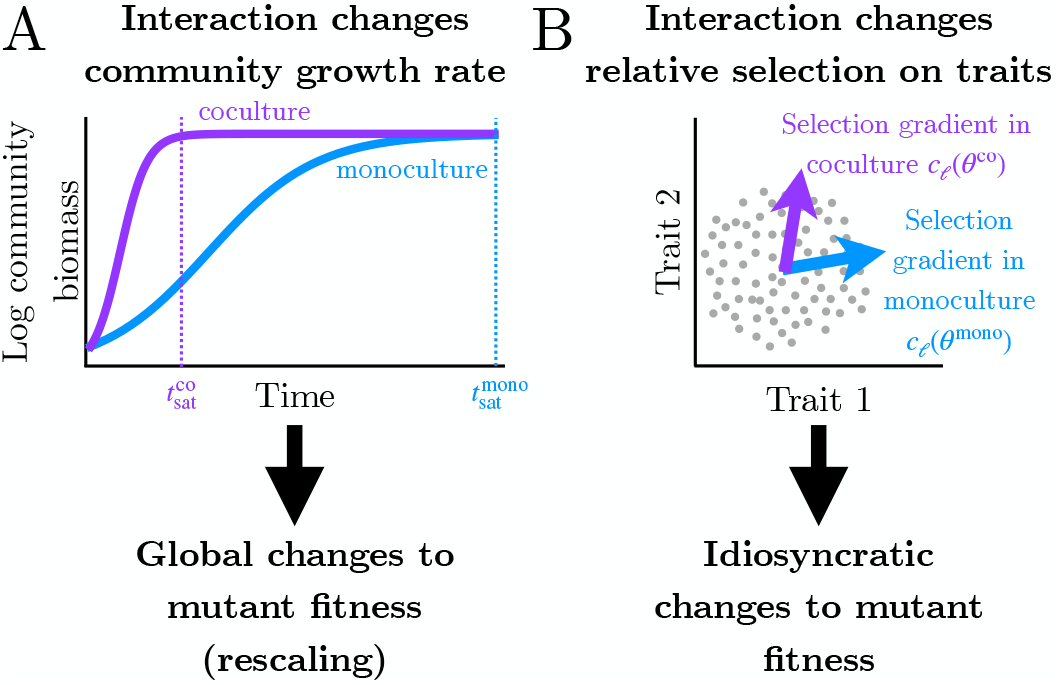
General mechanisms of how ecological interactions change the DFE. (A) Schematic growth curves of a whole community showing how an interaction can change its overall growth rate (∼1*/t*_sat_), leading to global changes in the DFE. (B) Schematic of how an interaction can change relative selection on traits, leading to idiosyncratic changes in the DFE.

Finally, we can determine the generality of idiosyncratic effects on the DFE by calculating the correlation coefficient between fitness in monoculture (*θ*^mono^) and in coculture (*θ*^co^). In Supplementary Sec. S5 we show that the correlation coefficient depends on the angle in trait space {*z*_𝓁_} between the selection gradients {*c*_𝓁_ (*θ*^mono^)} and {*c*_𝓁_ (*θ*^co^)}:

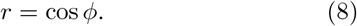

Hence, a change in the relative selection gradients *c*_*𝓁*_(*θ*) between traits will lead to a rotation in this space (Fig. 5B; also compare Fig. 4C to Fig. 4D), hence inducing idiosyncratic effects on fitness and reducing the correlation coefficient between monoculture and coculture. Any mechanism by which the interaction has disparate effects on selection for different traits will cause this, but one general mechanism for idiosyncratic effects is if mutations affect some traits which have effects that accrue per generation (and thus have selection gradients that change with the community growth rate) and other traits that do not.

Idiosyncratic effects therefore generally require pleiotropy (mutations affect multiple traits simultaneously); if mutations only affect one trait *z*_𝓁_, then fitness with *θ*^mono^ vs. *θ*^co^ will always be perfectly correlated because they are always proportional to each other by a factor of *c*_𝓁_ (*θ*^mono^)*/c*_𝓁_ (*θ*^co^). The exception to this occurs when the selection gradient for that one trait is exactly zero in one environment and nonzero in the other (e.g., a non-neutral trait in monoculture becomes neutral in the coculture); in that case, there is zero correlation between fitness in the two environments, and hence the interaction has maximum idiosyncratic effects without pleiotropy.

## DISCUSSION

### Interactions with faster or less efficient competitors globally reduce mutant fitness magnitude

Our model identifies change in the community growth rate as a primary mechanism by which competition changes mutant fitness, since fitness measured over a whole batch growth cycle is inversely proportional to community growth rate (Fig. 3C). Faster and slower competitors have asymmetric effects on the whole community’s growth rate, however (Fig. 3D), and thus faster or less efficient competitors globally reduce mutant fitness magnitude, while slower or more efficient competitors have little effect. The most direct test of our model comes from Martinson et al. [30], which found that competitive interactions with *E. coli* and *M. extorquens* had little effect on the DFE of *S. enterica*. In light of the model, this outcome is not surprising, since the growth rate of *S. enterica* is similar to that of *E. coli* and much faster than that of *M. extorquens*. However, the model also shows that this result was not inevitable: interactions of *S. enterica* with other competitor species (especially faster) could very well cause major DFE changes. A systematic test of the model predictions would be to take an ancestral species and its mutant library and then measure mutant fitness across cocultures with a range of other species. By measuring the growth rates of those communities, one can test the correlation of that community growth rate with the average magnitude of mutant fitness effect.

The model’s implication for evolution is that competition is likely to either slow the rate of adaptation (per batch growth cycle) or leave it unchanged, but it is unlikely to accelerate adaptation. This is consistent with some studies that find that interactions inhibit adaptation rate [33, 35] (although those communities involved a combination of cross-feeding and competition), but contrasts with another recent study that found that competition between species drove the evolution of stronger antibiotic resistance [37]. However, the latter may have been dictated by idiosyncratic changes to a small number of large-effect mutations, rather than the global change in the DFE.

An important caveat of these results is that the competitor must be significantly faster than the ancestral focal species to have a major impact on the DFE through these dynamics (e.g., Figs. 3B and 4B). Microbial growth rates do vary across orders of magnitude in nature [53, 54], although not necessarily between direct competitors. Coexistence between competitive species is possible under these dynamics [45, 46] as well as in other models of competition with a single resource [55, 56], but since those interactions do not change the community growth rate or the relative selection on traits (growth rate and lag time), they do not change the DFE. Thus the competitive interactions described here must be transient, since one species will eventually exclude the other under resource competition.

### There are two generic mechanisms of how interactions can change the DFE: change in community growth rate globally reshapes the DFE, while change in relative selection on traits causes idiosyncratic effects on mutant fitness

While the resource competition model presented here (Fig. 2) is intended to capture only the most basic features of microbial population dynamics and ecological interactions, it reveals these two mechanisms that can be defined generally in a model of arbitrary quantitative traits (Fig. 5). The two mechanisms correspond to the two types of statistical effects presented in Fig. 1 and show that any ecological interaction, including cooperative (e.g., crossfeeding [16–18]) and antagonistic (e.g., predator-prey [42] or secretion of toxins [40, 41]) interactions as well as competition [13–15], may be able to generate both effects. As a result, it is likely difficult if not impossible to infer an underlying interaction purely by considering its statistical effects on the DFE. This is also apparent in the empirical data of DFEs across systems, where the qualitative interaction type does not obviously dictate the statistical effects (Fig. 1C and Supplementary Fig. S3).

However, one can use the idiosyncratic effects of an interaction on specific mutations to deduce clues about the interaction [31]. For example, the idiosyncratic effects of a change in community growth rate can determine what mutations have effects that accrue per generation vs. per batch growth cycle. Li et al. [52] demonstrated this previously in yeast monocultures by artificially changing the length of batch growth cycles, but a similar effect should be revealed across multispecies communities as well. For example, mutations whose effects become weaker upon an increase in community growth rate are likely to have effects per generation, while mutations whose effects are independent of the interaction are likely to have effects accruing per batch growth cycle.

### Effect of interactions on mutant fitness suggests that higher-order interactions are pervasive in microbes

Since we can consider the fitness of a mutant relative to its ancestor as a type of interaction between them (competition), the ability of an interaction with another species to affect a mutant’s fitness is therefore an example of a higher-order ecological interaction [30, 46, 57– 60]. The pervasive evidence for ecological interactions between microbes affecting mutant fitness [23–31] implies that higher-order interactions, in this sense, are actually common among microbes, contrary to some studies that found limited evidence for them [30, 58]. This suggests that part of the challenge in finding higher-order interactions is how they are defined and tested. Most studies test them between species, whereas this argument means that they are more common within species (between a mutant and its ancestor).

## METHODS

### Analysis of empirical DFE data sets

We collected 36 pairs of coculture-monoculture DFE data sets (Supplementary Table S1, Supplementary Fig. S1) from 7 studies [23, 25, 27–31]. The mutants in all these studies are generated by transposon insertion mutagenesis, except for Venkataram et al. [28], which uses mutants spontaneously generated in an evolution experiment. We obtained all the fitness data from supplementary files published with the original papers. For data sets with replicate measurements, we averaged replicates for each mutant in each condition and calculated standard error of that mean, if those values were not already reported in the published data file. We excluded any mutants that had a missing fitness value in either the monoculture or coculture.

We calculated the following global effects of the interaction on the DFE: relative change in mean |(Δ*µ/µ*| = |(*µ*^co^ − *µ*^mono^) */µ*^mono^| , where *µ*^mono^ and *µ*^co^ are the DFE means in monoculture and coculture), relative change in standard deviation (|Δ*σ/σ*| = |(*σ*^co^− *σ*^mono^) */σ*^mono^| , where *σ*^mono^ and *σ*^co^ are the DFE standard deviations in monoculture and coculture), and earth mover’s distance (equivalent to 1-Wasserstein distance). Note that earth mover’s distance carries units of fitness and thus scales with the range of fitness measurements, which is larger in some data sets (e.g., Venkataram et al. [28]).

We measured the idiosyncratic effect of the interaction on the DFE as 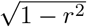, where *r* is the Pearson correlation coefficient between the monoculture and coculture fitness values (Supplementary Sec. S1). This definition is equivalent to taking the square root of the normalized sum of the square residuals from a linear regression (i.e., the deviations from the dashed line in Fig. 1B). To account for measurement noise in fitness, which might inflate the apparent idiosyncratic effect of the interaction (e.g., fitness in monoculture and in coculture might appear less correlated if measurements are noisier), we performed a maximum likelihood estimate for the true correlation between environments assuming Gaussian measurement noise with standard error on the mean taken from the replicate measurements (Supplementary Fig. S1). For all statistics we used standard functions in NumPy [61] or SciPy [62].

### Model of population dynamics and resource competition

We assume growth is limited by a single resource with concentration *R*(*t*), so that competition for this resource is the only interaction between genotypes. The biomass *N*_*k*_(*t*) of each species *k* and resource concentration *R*(*t*) change over time *t* according to

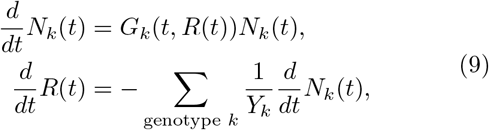

where *Y*_*k*_ is the yield of genotype *k* (biomass produced per unit resource; this assumes resources are consumed only for cell division and not for maintenance of existing biomass [63]) and *G*_*k*_(*t, R*(*t*)) is the per-capita growth rate of genotype *k* that depends on both time *t* and the available resource *R*(*t*). Note that the fundamental assumption of a consumer-resource model like this is that genotypes interact with each other only through the resource concentration *R*(*t*), since *G*_*k*_ does not depend on the other genotypes explicitly.

The growth rate function is defined as (Fig. 2)

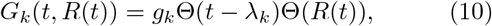

where Θ is the step function (Θ(*x*) = 0 if *x* < 0 and Θ(*x*) = 1 if *x* > 0). The saturation time *t*_sat_ when the resource is fully depleted and all growth stops is therefore defined by *R*(*t*_sat_) = 0. Substituting the growth rate function of Eq. 10 into Eq. 9 and integrating over time, we can obtain an implicit equation for *t*_sat_:

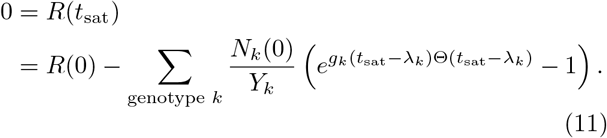

We define fitness of the mutant relative to its ancestor as [43, 45, 46]

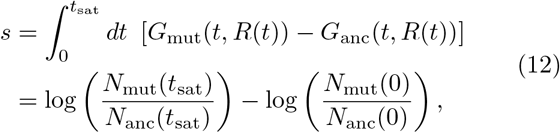

which in this specific model (Eq. 10) takes the form

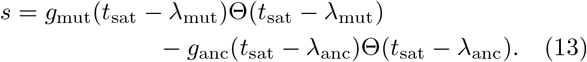

While it is possible to approximate (in the limit of weak selection, |*s*|≪ 1) an analytical form of the relative fitness showing its explicit dependence on the underlying growth traits [45, 46], for all figures in this article we numerically solve the saturation equation (Eq. 11) for *t*_sat_ and substitute that solution into Eq. 13 to calculate fitness.

### Simulation of mutations

We assume each mutant starts at a very low frequency (10^*−*6^) relative to its ancestor, to mimic a mutation spontaneously arising in a single cell. While we only consider one mutant at a time competing with its ancestor, we repeat this for a large number of mutants to estimate the DFE for all spontaneous mutations. In this model, the mutant genotype is characterized by three quantitative growth traits: its lag time *λ*_mut_, its maximum growth rate *g*_mut_, and its yield *Y*_mut_. We assume the ancestral genotype has trait values *λ*_anc_ = 1, *g*_anc_ = 1, and *Y*_anc_ = 1. We randomly sample trait values for 10^4^ mutants such that each trait value is drawn independently from a Gaussian distribution with the ancestral trait value as the mean and 1% of the ancestral trait value as the standard deviation. We assume mutations do not affect yields, since at low frequency their yields do not affect the overall population dynamics (their resource consumption is negligible when that rare). We assume the initial resource concentration is *R*(0) = 10^3^ and total initial biomass is Σ _genotype *k*_ *N*_*k*_(0) = 1 (so the fold-change of the monoculture is 10^3^). For monocultures of the ancestor and its mutant, this breaks down to *N*_anc_(0) = 1− 10^*−*6^ and *N*_mut_(0) = 10^*−*6^, while for the cocultures with the competitor, the initial biomasses are *N*_anc_(0) = 0.5 −5× 10^*−*7^, *N*_mut_(0) = 5× 10^*−*7^, and *N*_com_(0) = 0.5 (i.e., same total biomass and mutant frequency as in the monoculture but with 1:1 proportion between the competitor and the combined ancestor and its mutant). The traits of the competitor are varied as discussed in the main text.

## ACKNOWLEDGMENTS

NIH award R35GM16022 and the Human Frontier Science Program award RGEC30/2024 supported DGS and MM. We thank Jonathan Martinson for help re-analyzing his data.

## Notes

### Competing Interest Statement

The authors have declared no competing interest.

