## Supplementary Information for "A theoretical framework for how ecological interactions between microbes affect mutant fitness"

Justus Wilhelm Fink,[1,*] Duhita G. Sant,[1,*] and Michael Manhart[1,2,†]
[1] Center for Advanced Biotechnology and Medicine, Rutgers University, Piscataway, NJ, USA
[2] Department of Biochemistry and Molecular Biology, Robert Wood Johnson Medical School, Rutgers University, Piscataway, NJ, USA
(Dated: May 22, 2026)

### S1. GLOBAL AND IDIOSYNCRATIC EFFECTS WITH GAUSSIAN DFES

Here we provide an explicit mathematical example of global and idiosyncratic effects under the assumption that the distributions of $s_i^{\mathrm{mono}}$ and $s_i^{\mathrm{co}}$ across mutants are Gaussian, with means $\mu^{\mathrm{mono}}$ and $\mu^{\mathrm{co}}$ and standard deviations $\sigma^{\mathrm{mono}}$ and $\sigma^{\mathrm{co}}$. The Pearson correlation coefficient between $s_i^{\mathrm{mono}}$ and $s_i^{\mathrm{co}}$ is $r$. The coculture fitness of mutant $i$ is therefore related to its monoculture fitness according to:

$$s_i^{\mathrm{co}} = \underbrace{\mu^{\mathrm{co}} + \rho \frac{\sigma^{\mathrm{co}}}{\sigma^{\mathrm{mono}}}(s_i^{\mathrm{mono}} - \mu^{\mathrm{mono}})}_{\text{global effect } f(s_i^{\mathrm{mono}})} + \underbrace{\sigma^{\mathrm{co}}\sqrt{1-r^2}\eta_i}_{\text{idiosyncratic effect}} . \tag{S1}$$

In this case, we see that the global effect of the interaction is a linear transformation $f(s^{\mathrm{mono}})$ of monoculture fitness (shifting and rescaling according to the difference in moments between DFEs, independently of the individual mutant $i$; e.g., main text Fig. 1A). This global effect is what transforms the DFE's mean and standard deviation from $\mu^{\mathrm{mono}}$ and $\sigma^{\mathrm{mono}}$ in the monoculture to $\mu^{\mathrm{co}}$ and $\sigma^{\mathrm{co}}$ in the coculture. The idiosyncratic effect, however, is only nonzero if fitness is imperfectly correlated between environments ($|r| < 1$), in which case it depends on a random variable $\eta_i$ (drawn from a standardized Gaussian distribution, with zero mean and unit variance) that is unique to each mutant $i$ (e.g., main text Fig. 1B). Since $\sigma^{\mathrm{co}}$ defines the scale of fitness in the coculture and the random variable $\eta_i$ has unit variance, we take the factor $\sqrt{1-r^2}$ as a convenient dimensionless measure of the idiosyncratic effect, suitable for comparison across data sets where fitness may differ in scale (main text Methods, main text Fig. 1C and Fig. S3) as well as in the model (main text Fig. 4B,F and Fig. S6). Equation S1 generalizes to non-Gaussian distributions using the probability integral transform and Sklar's theorem [1], although in those cases the idiosyncratic effect $\eta_i$ will contribute non-additively to fitness.

* These authors contributed equally.

### S2. DFE MOMENTS IN THE COMPETITION MODEL

One way to assess global effects of an interaction on the DFE is to see how the interaction changes the DFE's moments. In the main text we establish the effect of an interaction on mutant fitness in the model must be mediated by the saturation time $t_{\mathrm{sat}}$. To calculate moments of the DFE in the model, it is therefore convenient to rewrite the definition of mutant fitness in the model (main text Eq. 13, Methods) with the dependence on $t_{\mathrm{sat}}$ isolated:

$$s = (g_{\mathrm{mut}} - g_{\mathrm{anc}})t_{\mathrm{sat}} - g_{\mathrm{mut}}\lambda_{\mathrm{mut}} + g_{\mathrm{anc}}\lambda_{\mathrm{anc}}. \tag{S2}$$

Here we ignore the case where the resource is depleted before either the mutant or ancestor exits lag phase (e.g., we ignore the step functions in main text Eq. 13), since that does not occur with the parameter ranges we use in the numerical calculations and simplifies our subsequent analytical calculations.

Since the mutant traits $g_{\mathrm{mut}}$ and $\lambda_{\mathrm{mut}}$ are modeled as random variables across all mutants (Methods), we can calculate moments of the DFE in terms of moments of those mutant trait distributions. The DFE mean is therefore

$$\langle s \rangle = (\langle g_{\mathrm{mut}} \rangle - g_{\mathrm{anc}})t_{\mathrm{sat}} - \langle g_{\mathrm{mut}}\lambda_{\mathrm{mut}} \rangle + g_{\mathrm{anc}}\lambda_{\mathrm{anc}}. \tag{S3}$$

In our numerical simulations, we assume mutants have the same traits as the ancestor on average ($\langle g_{\mathrm{mut}} \rangle = g_{\mathrm{anc}}$, $\langle \lambda_{\mathrm{mut}} \rangle = \lambda_{\mathrm{anc}}$) and the growth rates and lag times are uncorrelated ($\langle g_{\mathrm{mut}}\lambda_{\mathrm{mut}} \rangle = \langle g_{\mathrm{mut}} \rangle \langle \lambda_{\mathrm{mut}} \rangle$). In that case, the mean fitness is zero regardless of the competitor (no dependence on $t_{\mathrm{sat}}$), confirming our numerical observation in main text Fig. 3A. The DFE variance, on the other hand, is

$$\begin{aligned}\langle s^2 \rangle = &\langle (g_{\mathrm{mut}} - g_{\mathrm{anc}})^2 \rangle t_{\mathrm{sat}}^2 \\ &- 2\langle (g_{\mathrm{mut}} - g_{\mathrm{anc}})(g_{\mathrm{mut}}\lambda_{\mathrm{mut}} - g_{\mathrm{anc}}\lambda_{\mathrm{anc}}) \rangle t_{\mathrm{sat}} \\ &+ \langle (g_{\mathrm{mut}}\lambda_{\mathrm{mut}} - g_{\mathrm{anc}}\lambda_{\mathrm{anc}})^2 \rangle.\end{aligned} \tag{S4}$$

Again assuming mean mutant traits are the same as the ancestor's and that the traits are uncorrelated, the variance simplifies to main text Eq. 2.

### S3. EFFECT OF QUANTIFYING FITNESS PER GENERATION ON DFE STATISTICS

The effect of a competitive interaction globally magnifying fitness of all mutants by changing the community's growth rate (measured by $t_{\text{sat}}$) is dependent on measuring fitness over the whole batch growth cycle (main text Eq. 1), which is common in many experimental [2–4] and modeling studies [5, 6] and has convenient theoretical properties [7]. However, some studies (most prominently the Long-Term Evolution Experiment in *E. coli* [8]) normalize fitness over a batch growth cycle by the number of generations. How does this normalization alter the apparent effect of an interactionon the DFE? Since the definition of fitness uses per-capita growth rate and hence entails natural logarithms (main text Eq. 12), it is more convenient to measure generations by the number of $e$-fold increases in the population rather than doublings. For the ancestor the number of $e$-fold increases is $g_{\text{anc}}(t_{\text{sat}} - \lambda_{\text{anc}})$, and thus the variance in fitness per generation is (dividing main text Eq. 2 by $g_{\text{anc}}^2(t_{\text{sat}} - \lambda_{\text{anc}})^2$; Fig. S4B)

$$\begin{aligned}\langle s^2 \rangle_{\text{per generation}} = & \left( \frac{\langle g_{\text{mut}}^2 \rangle}{g_{\text{anc}}^2} - 1 \right) \\ & + \frac{\langle g_{\text{mut}}^2 \rangle}{g_{\text{anc}}^2} \frac{\left( \langle \lambda_{\text{mut}}^2 \rangle - \lambda_{\text{anc}}^2 \right)}{(t_{\text{sat}} - \lambda_{\text{anc}})^2}.\end{aligned} \tag{S5}$$

If mutations do not affect lag times ($\langle \lambda_{\text{mut}}^2 \rangle - \lambda_{\text{anc}}^2 = 0$), then the $t_{\text{sat}}$ dependence drops out. In this case, the interaction has no effect on fitness per generation, because the only effect of the interaction is to rescale time, which is cancelled out by the per generation normalization. However, if there is mutant lag variation, then $t_{\text{sat}}$ dependence remains and the interaction has an opposite effect on fitness per generation compared to fitness per cycle: a faster competitor (shorter $t_{\text{sat}}$) increases DFE width per generation while decreasing DFE width per cycle (compare Fig. S4B with main text Fig. 3C).

The explanation for these opposite outcomes is that the effect of a mutation on growth rate accrues every generation [4]; hence its effect over a whole batch growth cycle is proportional to the number of generations, which is cancelled out if we normalize fitness by that number. On the other hand, the effect of a mutation on lag accrues per cycle, since the lag phase only occurs once in that time; thus a lag mutation's effect on fitness is independent of the number of generations. Normalizing fitness per generation spreads that fitness effect of lag mutations over all generations during the cycle, so a shorter cycle (because of a faster competitor) concentrates that fitness effect in fewer generations and makes selection per generation stronger. Note that quantifying fitness per generation has no impact on idiosyncratic effects, since the change in time scale drops out of the correlation coefficient $r$.

### S4. DEPENDENCE OF SATURATION TIME ON COMPETITOR TRAITS

Previous work [5, 6] derived an approximate expression for the saturation time $t_{\text{sat}}$ in the model as a function of all genotypes' traits (lag times $\lambda_k$, growth rates $g_k$, and yields $Y_k$) and the initial conditions (initial resource concentration $R(0)$ and initial biomass concentrations $N_k(0)$). In the case of an ancestor and competitor, the approximate saturation time is

$$t_{\text{sat}} \approx \bar{\lambda} + \frac{1}{\bar{g}} \log \left( 1 + \frac{R(0)\bar{Y}}{N_{\text{anc}}(0) + N_{\text{com}}(0)} \right), \tag{S6}$$

where

$$\bar{\lambda} = \frac{x_{\text{anc}} g_{\text{anc}} \lambda_{\text{anc}} / Y_{\text{anc}} + x_{\text{com}} g_{\text{com}} \lambda_{\text{com}} / Y_{\text{com}}}{x_{\text{anc}} g_{\text{anc}} / Y_{\text{anc}} + x_{\text{com}} g_{\text{com}} / Y_{\text{com}}} \tag{S7}$$

is the effective community lag time,

$$\bar{g} = \frac{x_{\text{anc}} g_{\text{anc}} / Y_{\text{anc}} + x_{\text{com}} g_{\text{com}} / Y_{\text{com}}}{x_{\text{anc}} / Y_{\text{anc}} + x_{\text{com}} / Y_{\text{com}}} \tag{S8}$$

is the effective per-capita community growth rate,

$$\bar{Y} = \left( \frac{x_{\text{anc}}}{Y_{\text{anc}}} + \frac{x_{\text{com}}}{Y_{\text{com}}} \right)^{-1} \tag{S9}$$

is the effective community yield, and

$$\begin{aligned} x_{\text{anc}} &= \frac{N_{\text{anc}}(0)}{N_{\text{anc}}(0) + N_{\text{com}}(0)}, \\ x_{\text{com}} &= \frac{N_{\text{com}}(0)}{N_{\text{anc}}(0) + N_{\text{com}}(0)} \end{aligned} \tag{S10}$$

are the ancestor's and competitor's relative fractions of initial biomass. We neglect the mutant in these expressions because we assume its frequency is so low that it does not affect growth of the mutant or competitor (main text Methods). The salient feature of these equations is that $t_{\text{sat}}$ depends on $g_{\text{com}}$ through a reciprocal ($1/\bar{g}$), so that when $g_{\text{com}} < g_{\text{anc}}$ (slower competitor), $1/\bar{g}$ depends weakly on $g_{\text{com}}$, but when $g_{\text{com}} > g_{\text{anc}}$ (faster competitor), $1/\bar{g}$ depends strongly on $g_{\text{com}}$. This is why $t_{\text{sat}}$ is only slightly longer with a slower competitor but much shorter with a faster competitor (main text Fig. 3D).

### S5. QUANTITATIVE TRAITS MODEL

Here we define further details of the general model of quantitative traits in the main text. Let the growth of the ancestral species depend on a set of quantitative traits $\{z_\ell\}$. We assume that mutation effects on these traits are pleiotropic but uncorrelated; we can always choose such a parameterization of traits by performing a rotation in the trait space to find orthogonal, uncorrelated axes of variation given a distribution of spontaneous mutants (i.e., a principal component analysis).

Assuming the mutation only makes small relative perturbations to these traits, we can approximate its fitness as (main text Eq. 4)

$$s(\theta) \approx \sum_{\text{trait } \ell} c_\ell(\theta) \frac{\Delta z_\ell}{z_\ell}, \tag{S11}$$

by defining the selection gradient $c_\ell(\theta)$ on trait $\ell$ [9] as (using main text Eq. 12)

$$c_\ell(\theta) = z_\ell \int_0^{t_{\text{sat}}(\theta)} dt \ \frac{\partial G_{\text{anc}}(\theta)}{\partial z_\ell}. \tag{S12}$$

This holds for arbitrary population dynamics and ecological interactions; the growth rate function $G_{\text{anc}}$ can depend on any resources, the abundances of other genotypes, as well as external factors. In this general case, $t_{\text{sat}}$ represents whatever time scale growth occurs over in the batch culture. Both of these components may depend on the ecological interaction $\theta$. Equation S12 implies that any traits $z_\ell$ whose effects accrue per generation [4] (meaning that the integrand of Eq. S12 scales with $t$ at least as a constant to leading order) will have selection gradients proportional to that growth time scale, which should be inversely proportional to the growth rate of the whole community (main text Eq. 5).

In our competition model, the selection gradients are

$$\begin{aligned} c_{\text{lag}} &= \lambda_{\text{anc}} \int_0^{t_{\text{sat}}} dt \ \frac{\partial}{\partial \lambda_{\text{anc}}} \left( g_{\text{anc}} \Theta(t - \lambda_{\text{anc}}) \right) \\ &= -g_{\text{anc}} \lambda_{\text{anc}} \\ c_{\text{growth}} &= g_{\text{anc}} \int_0^{t_{\text{sat}}} dt \ \frac{\partial}{\partial g_{\text{anc}}} \left( g_{\text{anc}} \Theta(t - \lambda_{\text{anc}}) \right) \\ &= g_{\text{anc}}(t_{\text{sat}} - \lambda_{\text{anc}}), \end{aligned} \tag{S13}$$

so that the fitness of a mutation is

$$s \approx -g_{\text{anc}} \lambda_{\text{anc}} \frac{\Delta \lambda}{\lambda_{\text{anc}}} + g_{\text{anc}}(t_{\text{sat}} - \lambda_{\text{anc}}) \frac{\Delta g}{g_{\text{anc}}}, \tag{S14}$$

which matches the exact expression (Eq. S2) up to first order in the trait perturbations. The selection gradient for $g_{\text{anc}}$ is inversely proportional to the community growth rate ($1/t_{\text{sat}}$) because mutation effects on that trait are accrued per generation, while the selection gradient for the lag time $\lambda_{\text{anc}}$ is constant in the community growth rate because mutation effects on it are accrued only once per growth cycle [4].

To generalize our previous results on how an interaction globally changes the DFE, we show how the DFE moments depend on the interaction parameter $\theta$. The mean of the DFE is (main text Eq. 6, compare to Eq. S3)

$$\langle s(\theta) \rangle = \sum_{\text{trait } \ell} c_\ell(\theta) \frac{\langle \Delta z_\ell \rangle}{z_\ell}. \tag{S15}$$

Assuming the interaction changes at least one of the selection gradients $c_\ell(\theta)$, then the interaction will also generally change the DFE mean if the average mutation trait effects $\langle \Delta z_\ell \rangle$ are not zero for all traits $\ell$. This did not occur in our competition model only because we assumed the average mutation trait effects to be zero ($\langle g_{\text{mut}} \rangle = g_{\text{anc}}$, $\langle \lambda_{\text{mut}} \rangle = \lambda_{\text{anc}}$; see Fig. 3A). We can similarly calculate the variance (main text Eq. 7, compare to main text Eq. 2):

$$\begin{aligned} &\langle (s(\theta))^2 \rangle - \langle s(\theta) \rangle^2 = \\ &\qquad \sum_{\text{trait } \ell} (c_\ell(\theta))^2 \, \frac{\langle \Delta z_\ell^2 \rangle - \langle \Delta z_\ell \rangle^2}{z_\ell^2}, \end{aligned} \tag{S16}$$

where we have invoked the assumption that mutation effects on traits are uncorrelated from each other.

Finally, we can determine the generality of idiosyncratic effects on the DFE by calculating the correlation coefficient between fitness in monoculture ($\theta^{\text{mono}}$) and in coculture ($\theta^{\text{co}}$):

$$\begin{aligned} r &= \left( \sum_{\text{trait } \ell} c_\ell(\theta^{\text{mono}}) c_\ell(\theta^{\text{co}}) \frac{\langle \Delta z_\ell^2 \rangle - \langle \Delta z_\ell \rangle^2}{z_\ell^2} \right) / \\ &\quad \sqrt{\sum_{\text{trait } \ell} (c_\ell(\theta^{\text{mono}}))^2 \, \frac{\langle \Delta z_\ell^2 \rangle - \langle \Delta z_\ell \rangle^2}{z_\ell^2}} \times \\ &\quad \sqrt{\sum_{\text{trait } \ell} (c_\ell(\theta^{\text{co}}))^2 \, \frac{\langle \Delta z_\ell^2 \rangle - \langle \Delta z_\ell \rangle^2}{z_\ell^2}} \\ &= \frac{\mathbf{u}(\theta^{\text{mono}}) \cdot \mathbf{u}(\theta^{\text{co}})}{\|\mathbf{u}(\theta^{\text{mono}})\| \|\mathbf{u}(\theta^{\text{co}})\|} \\ &= \cos \phi, \end{aligned} \tag{S17}$$

where we have invoked the assumption that mutation effects on traits are uncorrelated from each other. The vector $\mathbf{u}(\theta) = \{c_\ell(\theta) \sqrt{\langle \Delta z_\ell^2 \rangle - \langle \Delta z_\ell \rangle^2} / z_\ell\}$ is an element-wise product of the selection gradient and the relative trait standard deviations.


[1] A. Sklar. Fonctions de répartition à N dimensions et leurs marges. *Annales de l'ISUP*, 8:229–231, 1959.

[2] S. Kryazhimskiy, D. P. Rice, E. R. Jerison, and M. M. Desai. Global epistasis makes adaptation predictable despite sequence-level stochasticity. *Science*, 344:1519–1522, 2014.

[3] B. H. Good, M. J. McDonald, J. E. Barrick, R. E. Lenski, and M. M. Desai. The dynamics of molecular evolution

over 60,000 generations. *Nature*, 551:45–50, 2017.

[4] Y. Li, S. Venkataram, A. Agarwala, B. Dunn, D. A. Petrov, G. Sherlock, and D. S. Fisher. Hidden complexity of yeast adaptation under simple evolutionary conditions. *Curr Biol*, 28:515–525.e6, 2018.

[5] M. Manhart, B. V. Adkar, and E. I. Shakhnovich. Tradeoffs between microbial growth phases lead to frequency-dependent and non-transitive selection. *Proc R Soc B*, 285:20172459, 2018.

[6] M. Manhart and E. I. Shakhnovich. Growth tradeoffs produce complex microbial communities on a single limiting resource. *Nat Commun*, 9:3214, 2018.

[7] J. W. Fink and M. Manhart. Quantifying microbial fitness in high-throughput experiments. *eLife*, 13:RP102635, 2024.

| Reference | Focal species | Other species | Putative interaction | Reps. |
|---|---|---|---|---|
| Ascensao et al. 2023 [10] | *Escherichia coli* LTEE L strain | *Escherichia coli* LTEE S strain | Cross-feeding | 2 |
| Ascensao et al. 2023 [10] | *Escherichia coli* LTEE S strain | *Escherichia coli* LTEE L strain | Cross-feeding | 2 |
| LaSarre et al. 2020 [11] | *Escherichia coli* | *Rhodopseudomonas palustris* Nx | Cross-feeding | 4 |
| LaSarre et al. 2020 [11] | *Escherichia coli* | *Rhodopseudomonas palustris* NxΔAmtB | Cross-feeding | 4 |
| Martinson et al. 2023 [12] | *Salmonella enterica* | *Escherichia coli* | Cross-feeding | 5 |
| Martinson et al. 2023 [12] | *Salmonella enterica* | *Methylorubrum extorquens* | Cross-feeding | 5 |
| Martinson et al. 2023 [12] | *Salmonella enterica* | *Escherichia coli* | Competition | 5 |
| Martinson et al. 2023 [12] | *Salmonella enterica* | *Methylorubrum extorquens* | Competition | 5 |
| Morin et al. 2018 [13] | *Escherichia coli* | *Hafnia alvei* | Unknown (day 1) | 1 |
| Morin et al. 2018 [13] | *Escherichia coli* | *Hafnia alvei* | Unknown (day 2) | 1 |
| Morin et al. 2018 [13] | *Escherichia coli* | *Hafnia alvei* | Unknown (day 3) | 1 |
| Morin et al. 2018 [13] | *Escherichia coli* | *Geotrichum candidum* | Unknown (day 1) | 1 |
| Morin et al. 2018 [13] | *Escherichia coli* | *Geotrichum candidum* | Unknown (day 2) | 1 |
| Morin et al. 2018 [13] | *Escherichia coli* | *Geotrichum candidum* | Unknown (day 3) | 1 |
| Morin et al. 2018 [13] | *Escherichia coli* | *Penicillium camemberti* | Unknown (day 1) | 1 |
| Morin et al. 2018 [13] | *Escherichia coli* | *Penicillium camemberti* | Unknown (day 2) | 1 |
| Morin et al. 2018 [13] | *Escherichia coli* | *Penicillium camemberti* | Unknown (day 3) | 1 |
| Pierce et al. 2021 [14] | *Escherichia coli* | *Candida* 135E | Unknown | 3 |
| Pierce et al. 2021 [14] | *Escherichia coli* | *Debaryomyces* 135B | Unknown | 3 |
| Pierce et al. 2021 [14] | *Escherichia coli* | *Penicillium* 12 | Unknown | 3 |
| Pierce et al. 2021 [14] | *Escherichia coli* | *Penicillium* SAM3 | Unknown | 3 |
| Pierce et al. 2021 [14] | *Escherichia coli* | *Penicillium* RS17 | Unknown | 3 |
| Pierce et al. 2021 [14] | *Escherichia coli* | *Scopulariopsis* JB370 | Unknown | 3 |
| Pierce et al. 2021 [14] | *Escherichia coli* | *Scopulariopsis* 165-5 | Unknown | 3 |
| Pierce et al. 2021 [14] | *Escherichia coli* | *Fusarium* 554A | Unknown | 3 |
| Pierce et al. 2021 [14] | *Pseudomonas psychrophila* | *Candida* 135E | Unknown | 3 |
| Pierce et al. 2021 [14] | *Pseudomonas psychrophila* | *Debaryomyces* 135B | Unknown | 3 |
| Pierce et al. 2021 [14] | *Pseudomonas psychrophila* | *Penicillium* 12 | Unknown | 3 |
| Pierce et al. 2021 [14] | *Pseudomonas psychrophila* | *Penicillium* SAM3 | Unknown | 3 |
| Pierce et al. 2021 [14] | *Pseudomonas psychrophila* | *Penicillium* RS17 | Unknown | 3 |
| Pierce et al. 2021 [14] | *Pseudomonas psychrophila* | *Scopulariopsis* JB370 | Unknown | 3 |
| Pierce et al. 2021 [14] | *Pseudomonas psychrophila* | *Scopulariopsis* 165-5 | Unknown | 3 |
| Pierce et al. 2021 [14] | *Pseudomonas psychrophila* | *Fusarium* 554A | Unknown | 3 |
| Schreier et al. 2023 [15] | *Ruegeria pomeroyi* DSS-3 | *Vibrio hepatarius* HF70 | Unknown | 4 |
| Schreier et al. 2023 [15] | *Ruegeria pomeroyi* DSS-3 | *Marivivens donghaensis* HF1 | Unknown | 4 |
| Venkataram et al. 2023 [16] | *Saccharomyces cerevisiae* | *Chlamydomonas reinhardtii* | Cross-feeding | 3 |

TABLE S1. **Summary of empirical data sets comparing microbial DFEs with and without an ecological interaction.**

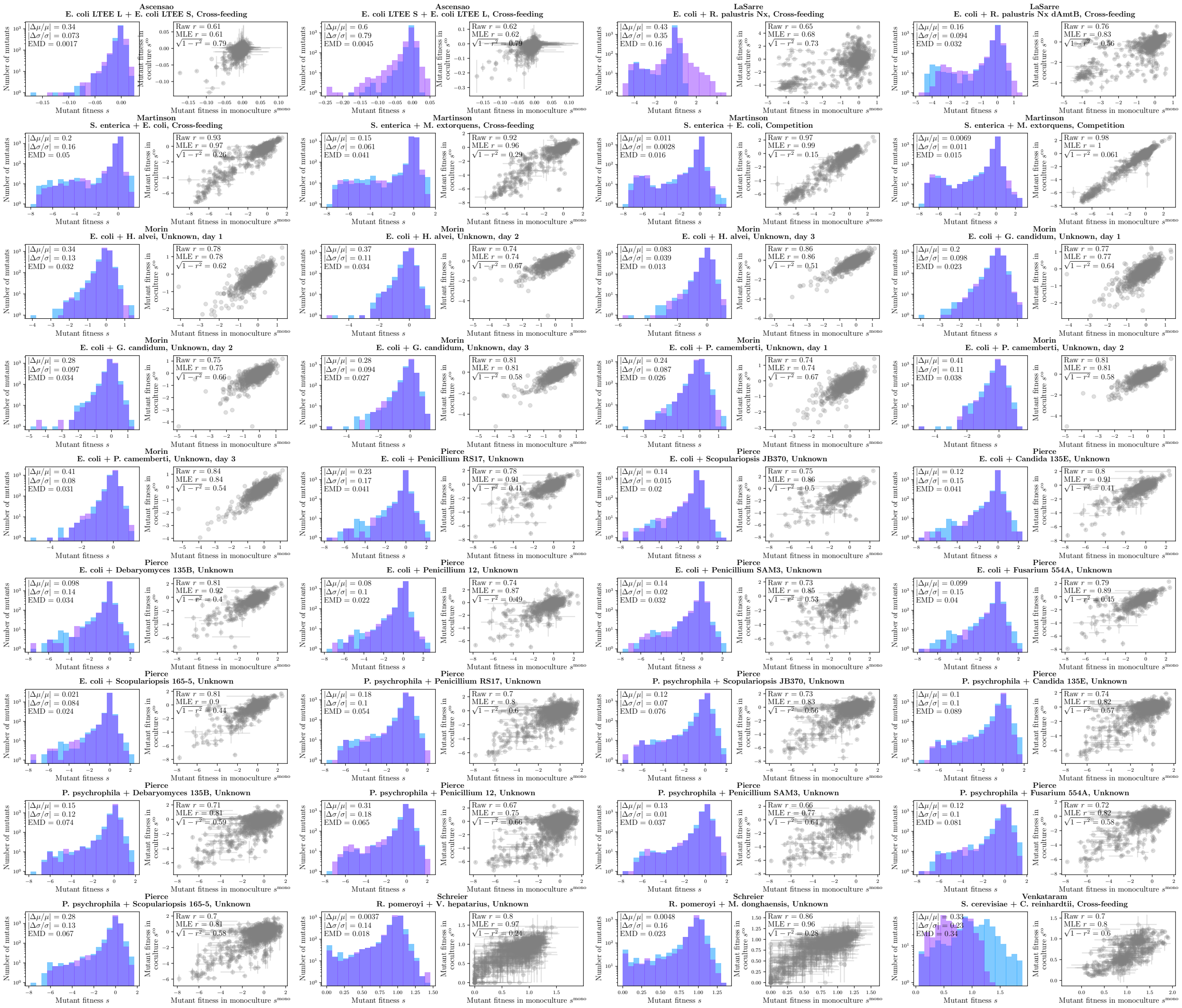


FIG. S1. **Empirical DFEs with and without ecological interactions.** Each pair of plots corresponds to a comparison of measured DFEs in monoculture and in a coculture with another strain or species. Above each pair of plots is the last name of the first author of the original study, and then the focal species (from which mutants are generated), the interacting species, and the putative interaction. See Table S1 for a complete list. The left plot in each pair shows histograms of the two DFEs (monoculture in blue, coculture in purple), with the calculated global effects printed in the upper left corner (main text Methods). The right plot shows a scatter plot of fitness for each mutant in monoculture vs. coculture, with error bars representing standard error on the mean over replicate measurements. The raw and maximum likelihood estimate of the Pearson correlation coefficient $r$ are printed in the upper left corner, along with the idiosyncratic effect (main text Methods).

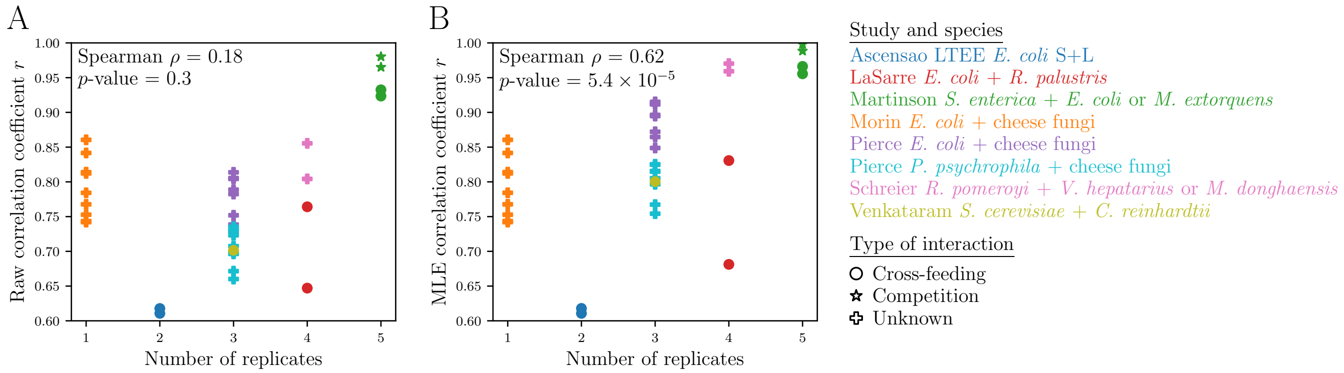


FIG. S2. **Dependence of number of replicates on apparent correlation between environments.** (A) Raw Pearson correlation coefficient $r$ of fitness between monoculture and coculture pairs (Table S1 and Fig. S1) as a function of the number of replicate measurements per mutant. (A) Same as (A) but using the maximum likelihood estimate of the Pearson correlation coefficient $r$ (main text Methods) to account for measurement noise. Each panel also shows the Spearman rank correlation $\rho$ and its associated $p$-value for the number of replicates vs. correlation coefficient $r$.

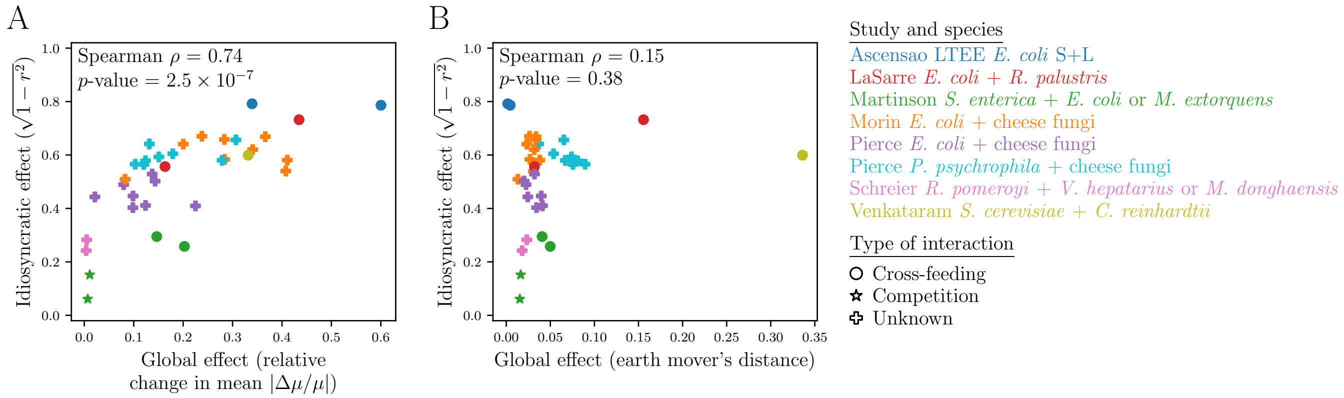


FIG. S3. **Comparing different measures of global effects on DFEs.** Same as main text Fig. 1C but measuring global effects using the (A) relative change in mean and (B) earth mover's distance (i.e., 1-Wasserstein distance; see main text Methods). Each panel also shows the Spearman rank correlation $\rho$ and its associated $p$-value for the global vs. idiosyncratic effects in that panel.

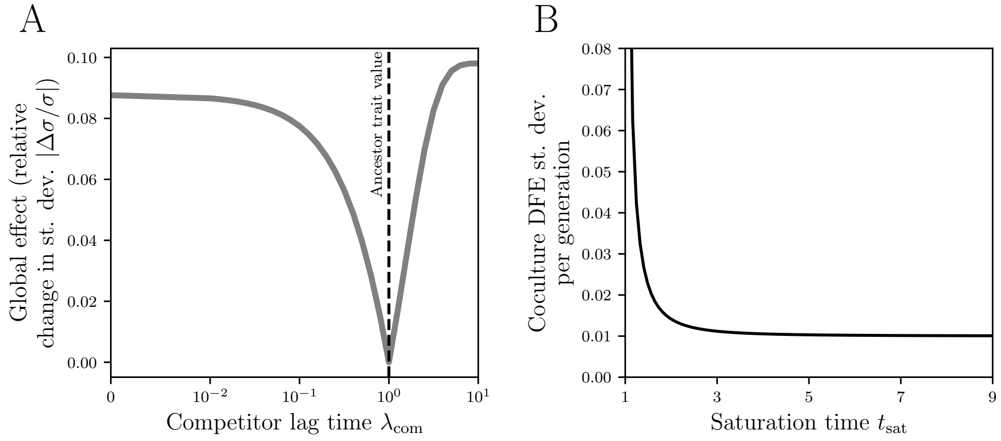


FIG. S4. **Effect of competitor lag time on DFE width and the effect of saturation time on fitness per generation.** (A) Same as main text Fig. 3B but as a function of competitor lag time $\lambda_{\mathrm{com}}$. (B) Dependence of DFE standard deviation per generation on saturation time $t_{\mathrm{sat}}$ (compare to main text Fig. 3C), based on the analytical calculation with fitness normalized per generation (Eq. S5).

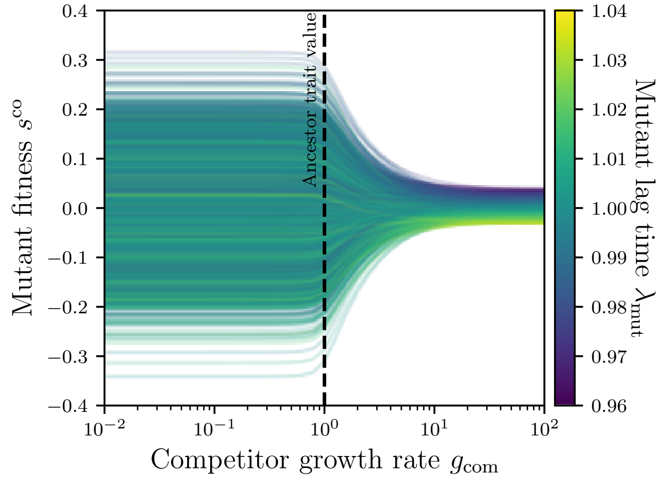


FIG. S5. **Effect of competitor growth rate on individual mutant fitness.** Fitness of each mutant in coculture $s^{\mathrm{co}}$ as a function of competitor growth rate $g_{\mathrm{com}}$; each mutant's line is colored according to its lag time $\lambda_{\mathrm{mut}}$.

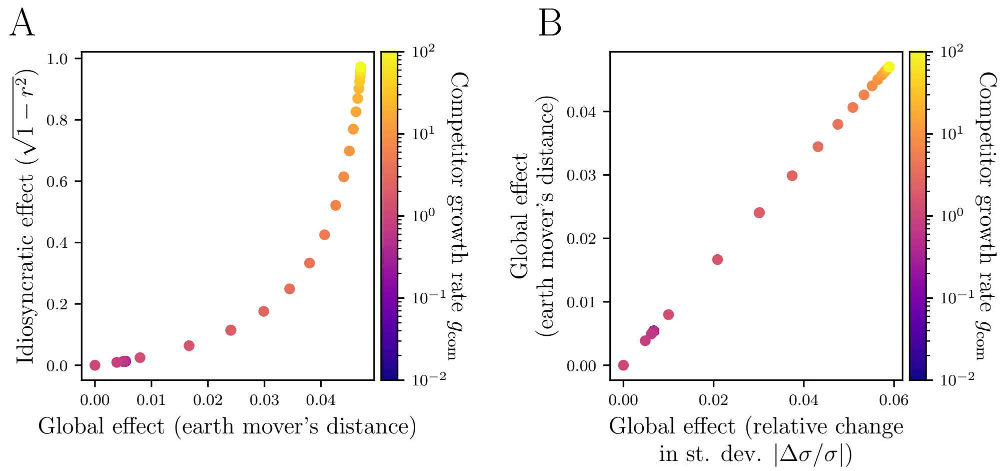


FIG. S6. **Global vs. idiosyncratic effects of competition.** (A) Same as main text Fig. 4F but calculating global effects as earth mover's distance (i.e., 1-Wasserstein distance; main text Methods) between the monoculture and coculture DFEs (compare to empirical data in Fig. S3B). (B) Comparison of global effects measured as relative change in DFE standard deviation vs. global effects measured as earth mover's distance. They are almost exactly proportional since the DFEs in the model are nearly Gaussian, and earth mover's distance = $\sqrt{2/\pi}\,|\sigma^{\mathrm{mono}} - \sigma^{\mathrm{co}}|$ for Gaussian distributions.
